# The Mitochondria-Modulating Small Molecule TEM1657 Suppresses NFKBIZ Pathway and Resolves Th17-Driven Psoriatic Inflammation

**DOI:** 10.64898/2026.10.03.756420

**Authors:** Preeti Gautam, Aleksander Salwinski, Ahmed B. Abdelwahab, Gaël Jalce, Laura Boulangé, Jessica Bettinger, Jordan Brouard, Alizée Etlicher, Romain Perouf, Julia Halper, Marie Millard, Natalia de Isla, Paul Hannewald, Jean-Francois Ginglinger, Marie-Thérèse Rubio, Jean-Yves Jouzeau, Frédéric Bourgaud, David Moulin

## Abstract

Psoriasis is a chronic skin disease characterized by red, scaly plaques mainly driven by dysregulated T cells that induce keratinocytes (KCs) proliferation and inflammation. Despite significant advancements in disease management through biologics and topical therapies, these treatments offer only temporary relief and fail to prevent disease recurrence. Considering the current therapeutic needs, we described herein a novel small molecule TEM1657, derived from 3,5-dicaffeoylquinic acid with anti-inflammatory potency in imiquimod-induced psoriasis mouse model as well as in Th17-stimulated KCs. Mechanistically, its action is mediated through modulation of mitochondrial dynamics, specifically via OPA1, an inner mitochondrial membrane protein that alters mitochondrial calcium influx, and further impacts cellular bioenergetics. Moreover, TEM1657 rapidly suppresses transcription of NFKBIZ and various other psoriasis-associated genes. These observations highlighted a pathway that represents an innovative target in psoriasis and other NFKBIZ-related disorders.

## Introduction

Psoriasis is a chronic autoimmune inflammatory disease characterized by the development of thick, scaly skin lesions that affect more than 125 million individuals worldwide^1,2^. Its pathogenesis arises from a complex interaction between abnormal keratinocyte (KCs) proliferation and immune responses, primarily mediated by T cells, leading to sustained inflammation and epidermal alterations^3^. The most prevalent clinical form of psoriasis is psoriasis vulgaris (plaque psoriasis), which accounts for 80-90% of psoriasis patients, and is represented by symptoms such as skin redness, itching and soreness^4,5^. Beyond skin manifestations, psoriasis is frequently associated with several comorbidities such as psoriatic arthritis, cardiometabolic disorders, obesity and hypertension, metabolic deficiencies, inflammatory bowel disease and psychiatric disturbances^6,7^. Despite extensive research, the precise etiology and pathophysiological mechanisms underlying psoriasis remain only partially understood^8^. Recent advances in immunopathology of disorder have, however, highlighted the key role of Th17 cells. Additionally, Th1 and Th22 cells also significantly contribute to maintaining the vicious cycle of inflammatory cytokines such as IL17, TNFα and IL23. The mechanistic insights gained in the last decade have transformed our understanding of psoriasis pathogenesis and paved the way for targeted biologic therapies that act on specific cytokine network ^9^. Presently, biologics such as anti-TNFα, infliximab and adalimumab and anti-IL17, secukinumab have been approved by both FDA and EMA^10,11^.

Another critical biomarker involved in the pathogenesis of psoriasis is Inhibitor of nuclear factor kappa B zeta (IκBζ) is an atypical coactivator of NFκB, encoded by NFKBIZ gene (nuclear factor of kappa light polypeptide gene enhancer in B cells inhibitor, zeta). Unlike other IκB proteins that inhibit NFκB in cytoplasm, IκBζ migrates to nucleus, where it binds to p50 subunit of NFκB and acts as a transcriptional coactivator of NFκB-responsive genes. In KCs, NFKBIZ acts as a direct transcriptional mediator of TNFα and IL17A stimuli, which consequently leads to transcription of psoriasis-associated genes^12,13^. Owing to its central role in psoriasis-specific inflammation, NFKBIZ emerges as a promising therapeutic target for localized inhibition to address a broad spectrum of patients with plaque psoriasis while minimizing potential systemic side effects^11,14^.

Caffeoylquinic acids (CQAs) are classified as phenolic specialized metabolites widely distributed across higher plants, including plant-based foods, with coffee, potatoes, and various vegetables being the main dietary sources. These molecules are derived from the phenylpropanoid biosynthesis pathway and share a common quinic acid (QA) core moiety esterified with one to four caffeoyl groups. CQAs are known to exhibit a wide range of biological activities such as antioxidant, anti-inflammatory, antibacterial, antiviral, and neuroprotective effects^15,16^. Among these, 3,5-dicaffeoylquinic acid (3,5-DiCQA, 1), also known as isochlorogenic acid A, has attracted particular attention for its anti-inflammatory properties^17^. This molecule was found to suppress TNF-α-and LPS-induced inflammation by downregulating IL-17 and MAPK signaling in both fibroblast-like synoviocytes and macrophages^18^. Moreover, it has been observed to attenuate inflammation-mediated pain hypersensitivity by promoting autophagy while suppressing MCP3-induced JAK2/STAT3 signaling in microglial cells^19^. Furthermore, 3,5-DiCQA (1) mitigates LPS-induced acute lung injury in mice by inhibiting NF-κB/NLRP3 signaling and reducing TNFα and IL-6 protein secretion in bronchoalveolar lavage fluid^20^.

In the present study, we describe TEM1657, a semisynthetic molecule derived from 3,5-dicaffeoylquinic acid, with anti-inflammatory properties that target a previously unexplored signaling axis by suppressing NFKBIZ’s transcription and potentially modulating OPA1 and mitochondrial homeostasis. Our findings identify a novel therapeutic axis and position TEM1657 as a promising therapeutic candidate for Th17-driven inflammatory disorders.

## Results

### Synthesis of TEM1657 an analogue of 3,5-DiCQA with enhanced anti-inflammatory potency

Building on previous studies demonstrating the anti-inflammatory properties of the natural compound 3,5-DiCQA^18,19^, we undertook a rational drug design approach to synthesize analogues optimized for both potency and topical administration in chronic inflammatory skin disease models^21^. The most promising candidate was identified through the replacement of the carboxylic acid moiety with an amide group; a modification specifically aimed at enhancing both anti-inflammatory efficacy and skin permeability. Herein, the amide analogue was synthesized in one step from 3,5-DiCQA through a peptide coupling reaction with the corresponding amine in presence of HOBt and EDCI (Fig. 1a). Further, the newly synthesized analogue, TEM1657 and its parent molecule 3,5-DiCQA were screened in an *in vitro* experiment for their ability to inhibit the secretion of inflammatory markers in human PMA challenged-keratinocytes. Interestingly, our data revealed that TEM1657 has greater potency than 3,5-DiCQA in suppressing pro-inflammatory cytokines IL-8, IL-6, TNF-α, PGE₂ and LTB₄ (Fig. 1b-f). Both compounds were further evaluated for their ability to inhibit NFKBIZ transcription in Th17-stimulated KCs. Notably, at a low dose (6.25 µM), only TEM1657 significantly downregulated NFKBIZ transcriptional expression (Fig. 1g). This enhanced activity may be attributed, at least in part, to the improved cellular permeability of TEM1657, which is more lipophilic and neutral at physiological pH than the natural parent molecule, 3,5-DiCQA. These observations clearly directed us to extend our studies on TEM1657 and explore the anti-inflammatory potential of this newly discovered molecule.

**Figure 1.**
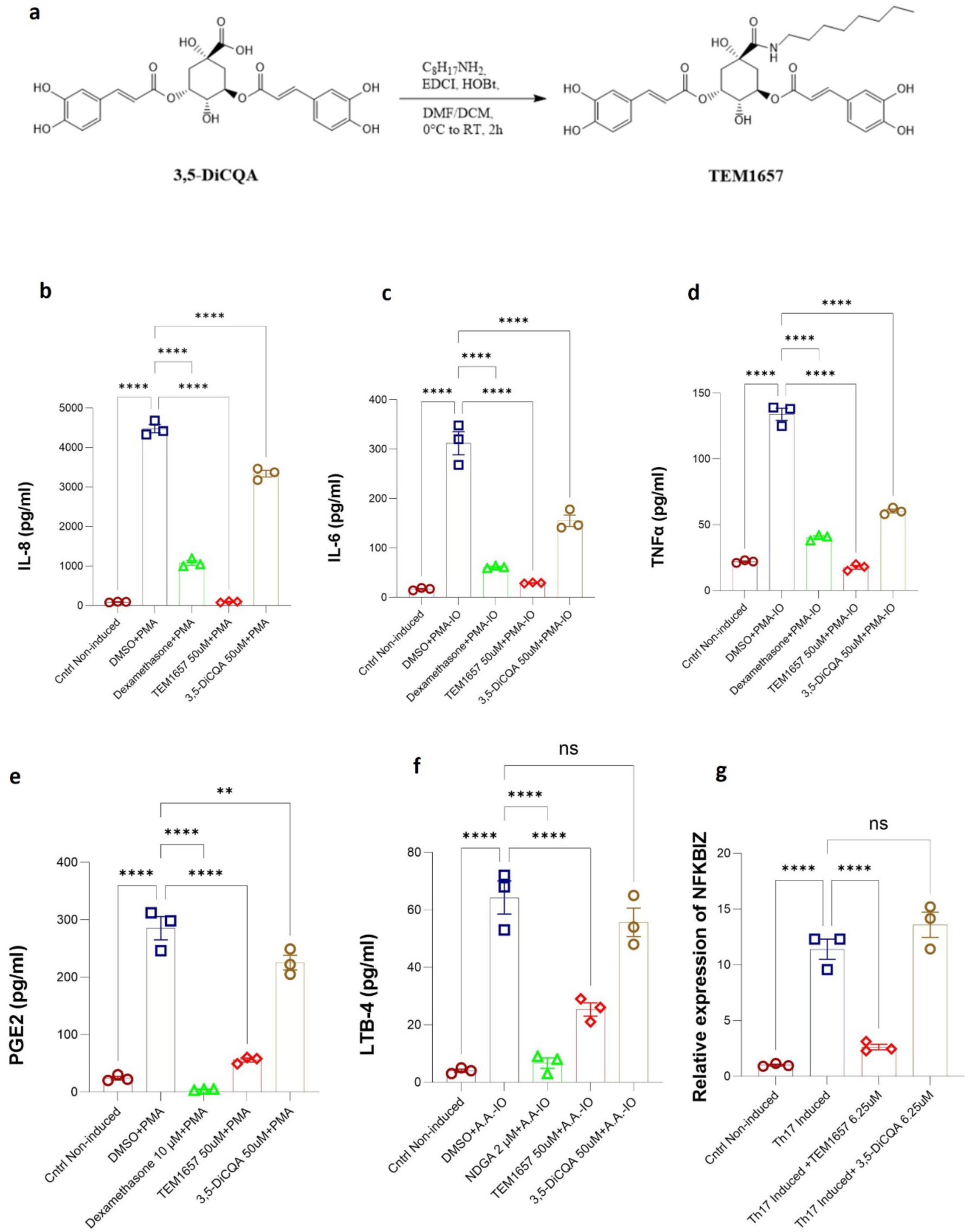
(a) Synthesis of TEM1657 from 3,5-DiCQA; concentrations of (b) IL-8 (c) IL-6 (d) TNFα (e) PGE2 (f) LTB-4, in the supernatant of KCs after incubation with TEM1657 and 3,5-DiCQA; (g) relative expression of NFKBIZ. The data are presented as the means ± standard errors of the means (S.E.M.) Statistical analysis was conducted by one way ANOVA and multiple comparisons were performed with Dunnett’s multiple comparisons test. *p < 0.05, **p < 0.01, ***p < 0.001, ****p < 0.0001, ns not significant.

### TEM1657 effectively alleviates psoriasis-like symptoms in the imiquimod-induced psoriasis mouse model

To evaluate TEM1657’s therapeutic efficacy, we employed the imiquimod (IMQ)-induced mouse model of psoriasis, wherein psoriatic lesions were topically induced over 11 days. The experimental design included multiple groups: TEM1657 administered at different doses, a naive control group, a therapeutic positive control (IMQ + Dermoval®, clobetasol propionate 0.05% cream), and a vehicle-induced control group (IMQ alone) to confirm successful disease induction^22,23^. Treatments were administered from day 7 to day 11 (Fig. 2a). Psoriasis-like skin manifestations (Fig.2b)—desquamation (Fig. 2d), erythema (Fig. 2e), and induration (Fig. 2f)— as well as the overall PASI score (Fig. 2g) showed significant dose-dependent improvement with TEM1657 compared to the IMQ group. Additionally, body weight (Fig. 2c) improved with the 0.8% dose, and all tested concentrations of TEM1657 reduced spleen size (Fig. 2h). These results confirm that in the IMQ-induced model known to reliably reproduce human psoriasis symptoms, TEM1657 attenuates disease manifestations in a dose-dependent manner.

**Figure 2.**
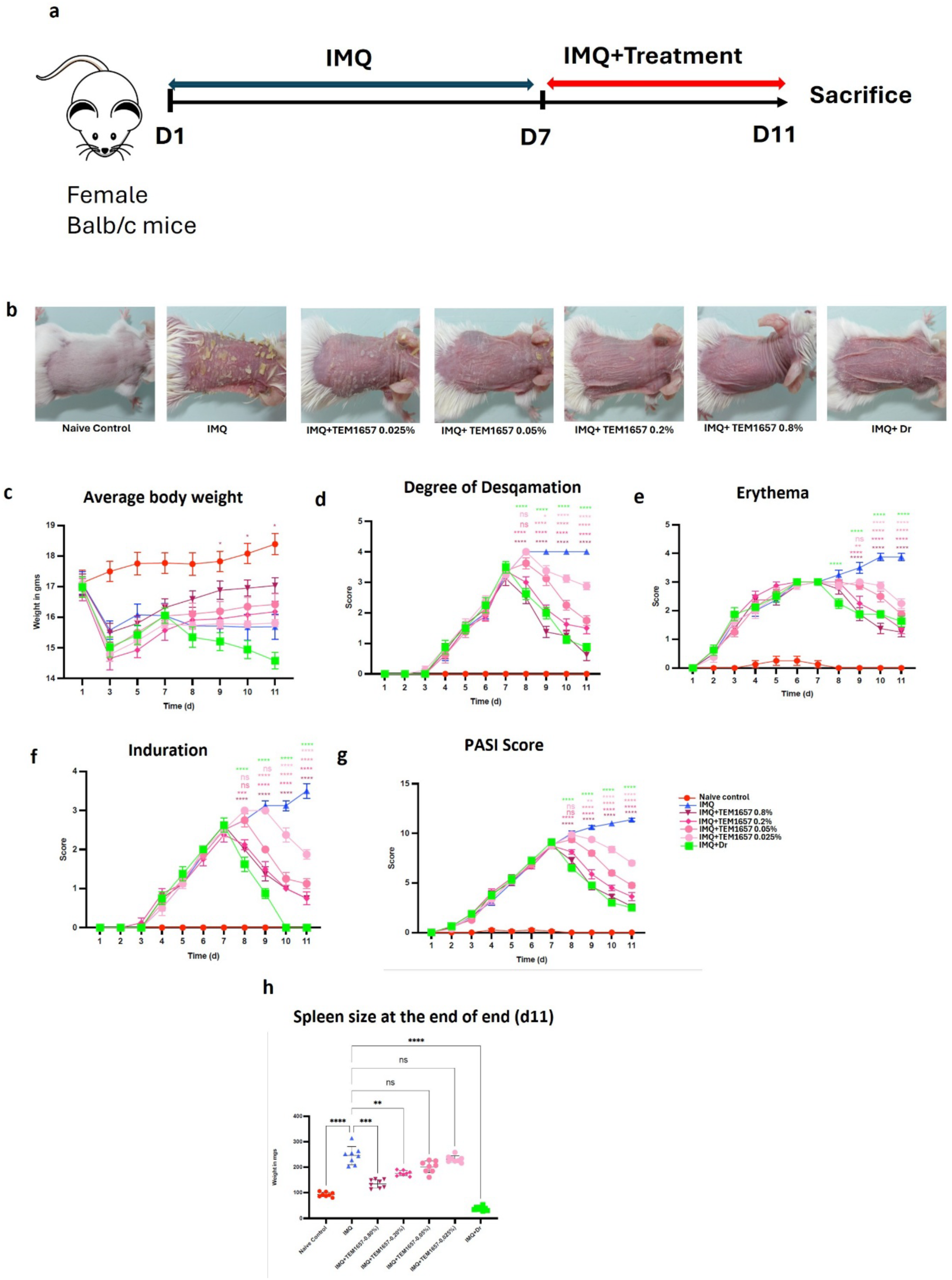
TEM1657 improves psoriasis-like symptoms in the imiquimod-induced psoriasis mouse model. (a). Experimental design (b). Macroscopic view of back skin. (c). Average body weight (d) Degree of desquamation (e) Erythema (f) Induration score (g) PASI score; of control (naive control), diseased (IMQ), IMQ + TEM1657 0.025-0.8% and IMQ+ Dermoval treated groups measured during the 11 days of experiment. (h) Size of spleen measured at the end of experiment. The data are presented as the means ± standard errors of the means (S.E.M.). All statistical analysis was conducted in the comparison to diseased (IMQ) group. Significance was identified with Two-way ANOVA and multiple comparisons were performed with Dunnett’s multiple comparisons test. *p < 0.05, **p < 0.01, ***p < 0.001, ****p < 0.0001, ns not significant

### PK/PD studies revealed that TEM1657 is distributed in blood cells within an hour and reaches the keratinocyte nucleus within 90 minutes

Mass spectrometry Single Reaction Monitoring (SRM) was used to investigate the distribution kinetics of TEM1657 in blood cells, plasma and different cellular compartments of KCs. Interestingly, we observed that within one hour, TEM1657 was cleared from plasma and entered blood cells (Sup Fig. S1a). Further analysis of its subcellular distribution in KCs showed that TEM1657 predominantly accumulated in the membrane fraction within 30 min (Supplementary Fig. S1b). Over the following 90 min, TEM1657 levels increased in the cytoplasmic fraction (Supplementary Fig. S1c), followed by its detection in the nuclear fraction (Supplementary Fig. S1d) and, ultimately, in chromatin-bound nuclear extracts (Supplementary Fig. S1e). This sequential localization pathway—membrane → cytoplasm → nucleus → chromatin—was consistent with high cell permeability and suggests potential nuclear activity of TEM1657.

### TEM1657 diminishes Th17-induced expression of major psoriasis-associated genes in KCs

We then explored the mechanism of action of this molecule. As synergistic stimulation with IL-17A and TNFα were known to upregulate the psoriasis signature genes^24^, we performed a transcriptomic analysis on KCs stimulated with Th17 cytokines (IL-17A and TNFα) for 90 min in the presence and absence of TEM1657^25^. The model’s functionality was validated by the expected overexpression of key proinflammatory cytokines and chemokines such as TNFα, CXCL8, CXCL2, CCL20, IL1β and IL36γ under Th17 stimulation, mimicking psoriasis-like inflammation^26^. Importantly, TEM1657 treatment significantly downregulated expression of major psoriasis-associated proinflammatory genes (Fig. 3a). Consistently, gene enrichment analysis^27^ showed that the overexpressed genes in the Th17 condition were highly enriched in TNFα and IL-17 signaling pathways as well as in the rheumatoid arthritis pathway(Fig. 3b). Interestingly, we found that many of the genes differentially expressed following TEM1657 treatment were associated with the same inflammatory pathways. Taken together, these observations strongly validate the biological activity of TEM1657 in mitigating Th17-driven inflammatory responses in KCs, thereby supports its therapeutic potential for psoriasis and other Th17-related inflammatory conditions.

**Figure 3.**
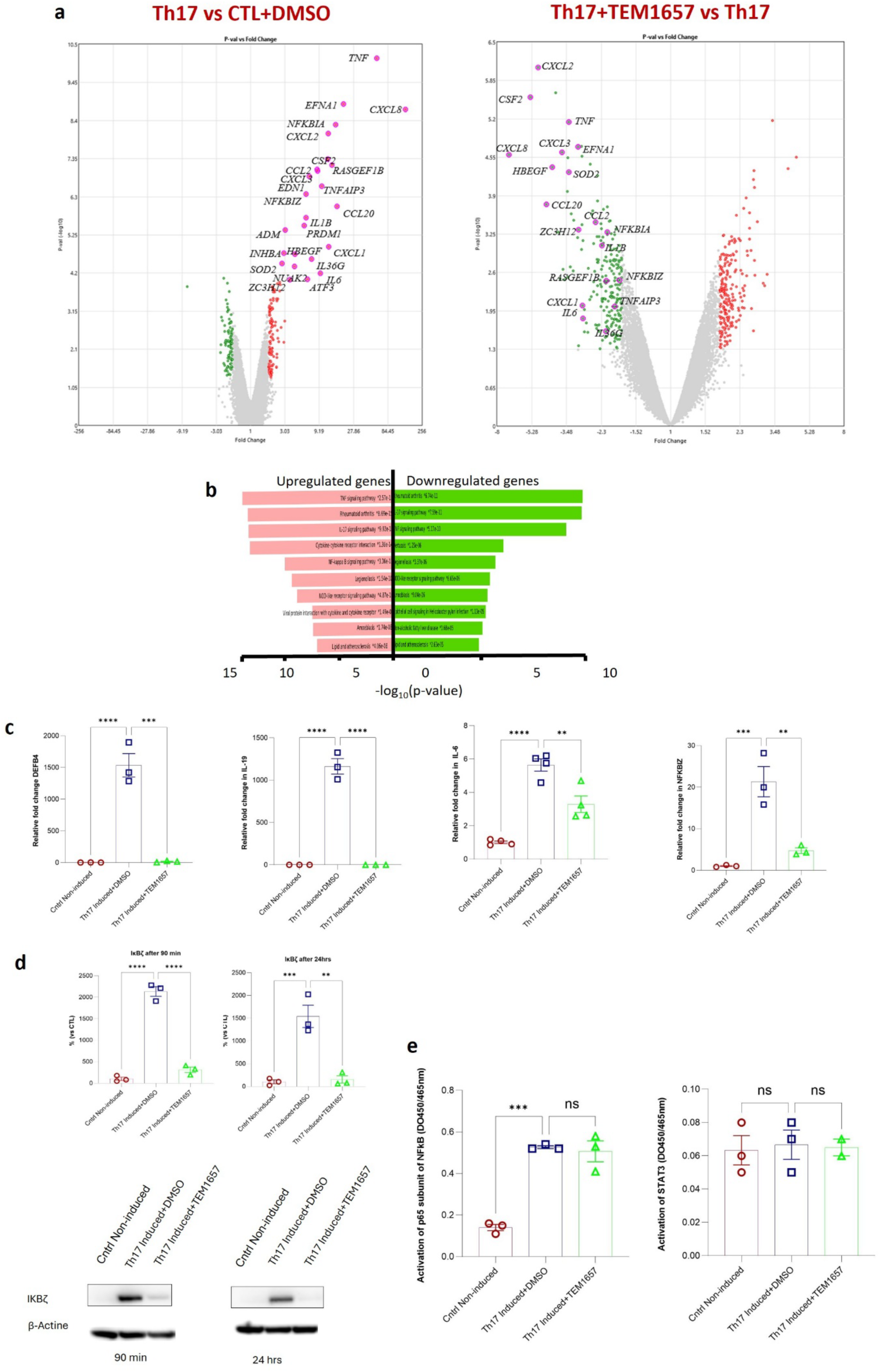
TEM1657 diminishes Th17-induced expression of major psoriasis-associated genes in KCs. (a) Microarray analysis for whole transcriptome profiling, volcano plot on right KCs with DMSO in comparison to Th17-induced KCs; volcano plot on left KCs with Th17 induced KCs+ TEM1657 in comparison to Th17-induced KCs. (b) Gene enrichment analysis of microarray genes (c) The relative expression of psoriasis-associated genes DEFB4 (after 24 hours), IL-19 (after 24 hours), IL-6 (after 90 minutes) and NFKBIZ (after 90 minutes). (d) Protein content of IκBζ (protein encoded by NFKBIZ) after 90 min and 24 hours incubation with TEM1657 in Th17-induced KCs; Lower panel, a representative image of protein expression analyzed with the help of western blot. (e) Trans AM experiment to analyze the activation of p65 subunit of NFκB and STAT3 after incubating Th17-induced KCs with TEM1657 for 6 hours. TEM1657 was used at a concentration of 6.25 μM in these experiments. The data are presented as the means ± standard errors of the means (S.E.M.) Statistical analysis was conducted by one way ANOVA and multiple comparisons were conducted with the help of Dunnett’s multiple comparisons test. *p < 0.05, **p < 0.01, ***p < 0.001, ****p < 0.0001, ns not significant

### TEM1657 inhibits Th17-induced expression of NFKBIZ

Tofurther validate the role of TEM1657, we quantified the mRNA expression of psoriasis-associated genes. Consistent with the earlier global transcriptomic analysis, it was observed that TEM1657 treatment significantly downregulated DEFB4, NFKBIZ, IL6, and IL19 in KCs^26^(Fig. 3c). Given the previously established role of NFKBIZ in psoriasis pathogenesis and inflammatory signaling^28,29^, we wanted to further investigate the impact of TEM1657 on its expression. Strikingly, we found that TEM1657 induced a rapid reduction in NFKBIZ transcriptional expression within 75–90 minutes of treatment (Sup Fig. S2). In line with the transcriptional expression, the protein levels of IκBζ (protein translated by mRNA NFKBIZ) were also found to be suppressed by TEM1657 in Th17-stimulated KCs (Fig. 3d). Together, these results identify TEM1657 as a potent modulator of keratinocyte-driven inflammation, acting through NFKBIZ/IκBζ repression and downstream cytokine inhibition, thereby providing a strong mechanistic rationale for its therapeutic potential in psoriasis.

### TEM1657 acts independently of NF-κB and STAT3

We further decided to investigate whether TEM1657 affects NF-κB and STAT3 activity. Given the established role of NFKBIZ/IκBζ in modulating NF-κB signaling, we sought to determine whether TEM1657-mediated suppression of the NFKBIZ/IκBζ axis was associated with altered NF-κB activation ^30^. In addition, STAT3 has been identified as a transcriptional activator of NFKBIZ, as it increases its transcription by directly binding to its promoter^31^. We now wondered if TEM1657 exerts its effect by modulating the activation of these two important transcription factors. To address this question, Trans AM assays were performed to assess NF-κB p65 or STAT3 activation in Th17-stimulated KCs treated with TEM1657 s^32^. We observed that TEM1657 had no detectable effect on the activation of either NF-κB p65 or STAT3 (Fig. 3e). These findings indicate that the anti-inflammatory effects of TEM1657 occur independently of canonical NF-κB pathway suppression. Importantly, sparing normal NF-κB’s physiological activity, which is essential for immune defense and cell survival. This observation recommends an additional therapeutic advantage of TEM1657, as broad NF-κB inhibition is frequently associated with adverse effects^33^. Thus, TEM1657’s selective mechanism, centered on NFKBIZ/IκBζ suppression without perturbing core NF-κB functions, highlights its potential as a promising and safer anti-inflammatory candidate for psoriasis and Th17-induced pathological conditions.

### CETSA® analysis revealed a mitochondrial-related mechanism of action for TEM1657

To investigate the mechanism of action of TEM1657, we employed the cellular thermal shift assay (CETSA®) in Th17-stimulated KCs with TEM1657. CETSA® is an unbiased, cell-based approach for target identification that detects compound-induced changes in protein thermal stability, thereby providing evidence of potential compound–protein interactions^34^. It was observed that TEM1657 altered the thermal stability of MGLL (monoglyceride lipase), OPA1 (optic atrophy protein 1), MICU1 (mitochondrial calcium uptake 1) and FECH (Ferrochelatase) (Fig. 4a). Herein, MICU1, OPA1 and FECH are inner mitochondrial membrane proteins, where OPA1 and MICU1 stabilize cristae formation and play a crucial role in unified mitochondrial calcium influx^35–37^. Interestingly, a recent mechanistic study showed that OPA1 also regulates mitochondrial calcium uptake through ER-mitochondrial coupling^38^.

**Figure 4.**
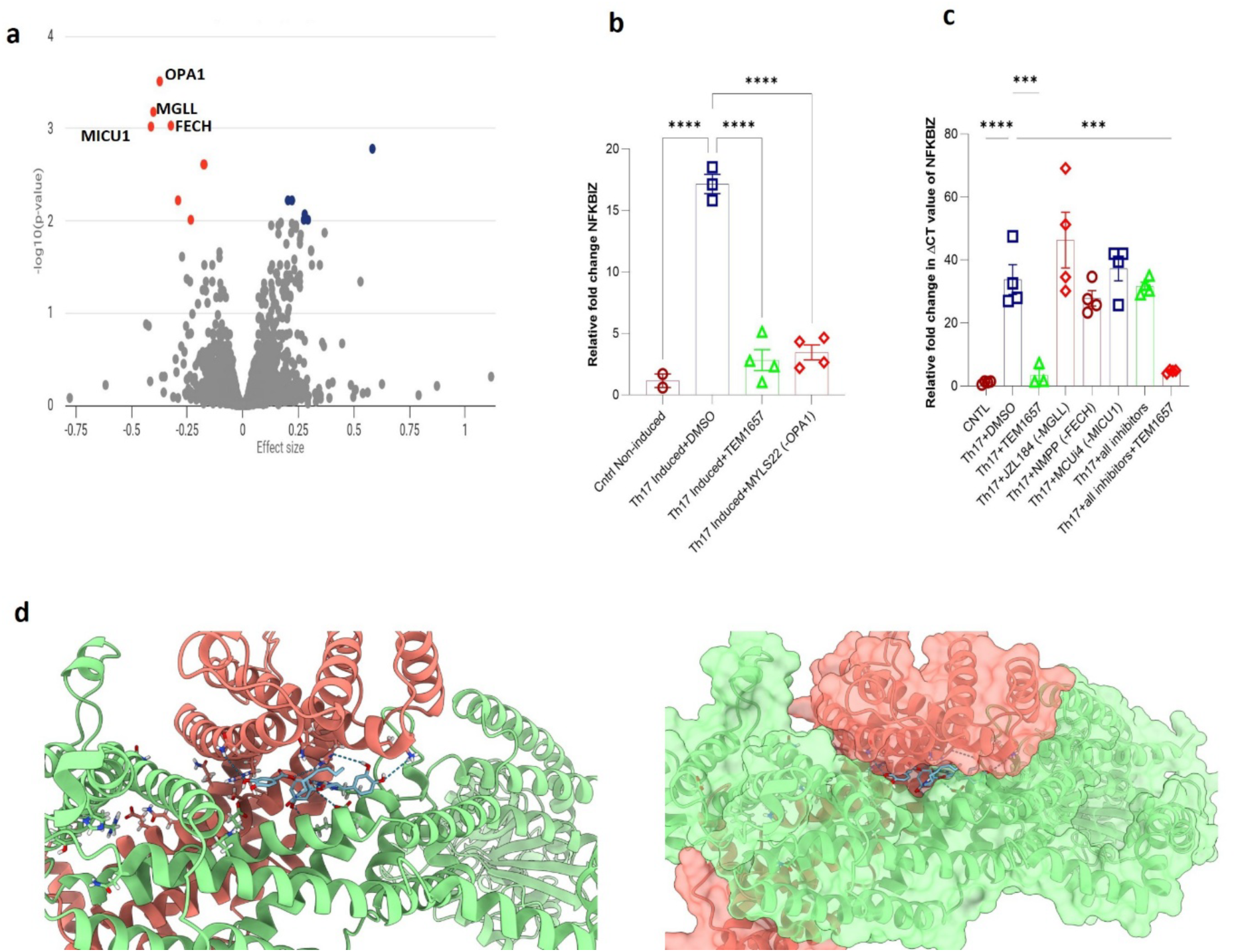
OPA1 might be the potential drug target for TEM1657 (a) Unbiased cellular target identification assay, CETSA® was performed after stimulating KCs with Th17-induction for 15 minutes, followed by 30 minutes incubation with TEM1657 (10μM). OPA1 emerged as a potential target in this assay (b) Relative expression of NFKBIZ in the presence of chemical inhibitor of OPA1-MYLS22. (c) Relative expression of NFKBIZ in the presence of chemical inhibitors of MGLL-JZL184, FECH-NMPP and MICU1-MCUi4. (d) The binding mode of one TEM1657 pose forming the maximum number of hydrogen bonds at the interface of two monomers. Each monomer was depicted as a transparent surface and a cartoon of a different color (light green and salmon). The data are presented as the means ± standard errors of the means (S.E.M.) Statistical analysis was conducted by one way ANOVA and multiple comparisons were conducted with the help of Dunnett’s multiple comparisons test. ***p < 0.001, ****p < 0.0001.

### OPA1: The potential mediator of TEM1657

Now a lingering question is whether the protein interactions in identified by CETSA® has functional relevance linked to NFKBIZ pathway and whether the inhibition of these proteins has any impact on transcriptional expression of NFKBIZ. To test this hypothesis, MGLL, OPA1, MICU1 and FECH were pharmacologically inhibited using their particular small-molecule inhibitors, JZL184, MYLS22, MCU-i4 and NMPP. OPA1 inhibition with MYLS22 markedly reduced NFKBIZ mRNA levels (Fig. 4b), whereas inhibition of MGLL, MICU1 and NMPP did not significantly alter NFKBIZ expression (Fig. 4c), indicating that, under these experimental conditions, OPA1 is the most likely of the CETSA®-identified candidates to be functionally linked to NFKBIZ regulation. OPA1 is a well-characterized mitochondrial protein localized to the inner mitochondrial membrane, where it regulates calcium influx and mitochondrial dynamics, thereby modulating cellular stress responses. This observation aligns with a recent study which observed that silencing OPA1 leads to decreased expression of multiple NF-κB inhibitors including NFKBIZ in vascular endothelial cells HUVECs^39^. The findings imply that OPA1 inhibition phenocopied TEM1657’s suppression of NFKBIZ and this finding strongly supports the conclusion that OPA1 is a functional mediator of TEM1657 activity. Furthermore, a docking simulation study using AUTODOCK VINA revealed that TEM1657 may cause interference in OPA1 oligomerization at the membrane-associated interface (Fig. 4d). Taken together, these findings support a model in which TEM1657 exerts its anti-inflammatory effect by modulating OPA1 function, which in turn may regulates NFKBIZ/IκBζ transcriptional activity. Consequently, we identified for the first time an unrecognized link between TEM1657, mitochondrial dynamics, and the NFKBIZ/IκBζ inflammatory axis.

### TEM1657 regulates mitochondrial calcium homeostasis

Guided by the crucial involvement of OPA1 in regulation of mitochondrial homeostasis by facilitating the transport of calcium ions from the cytosol into mitochondria, thereby modulating cytosolic calcium concentrations^40^, we wanted to investigate whether TEM1657 has any impact on real-time migration of calcium ions from cytosol to mitochondria. To address this question, we treated KCs with TEM1657 and stained them with cytosolic and mitochondrial Ca^2+^ indicators. Additionally, calcium mobilization was induced with the calcium ionophore ionomycin. Our flow cytometry data revealed that in Th17-induced KCs pretreated with TEM1657, both cytosolic and mitochondrial calcium concentrations were markedly reduced following ionomycin stimulation, whereas MYLS22 treatment had no detectable effect (Fig. 5a, b). These findings were additionally validated with the help of time-lapse confocal live-cell imaging. Consistently, TEM1657 treatment reduced the influx of calcium from the cytoplasm into mitochondria, while cytoplasmic calcium levels remained unchanged (Fig. 5c, d). Taken together, our results demonstrate that TEM1657 disrupts calcium transfer from cytosol to mitochondria, thus altering mitochondrial homeostasis. Importantly, this mitochondrial dysregulation via OPA1 translates into extra-mitochondrial effects, where TEM1657 modulates IκBζ/NFKBIZ expression independently of canonical NF-κB signaling.

**Figure 5.**
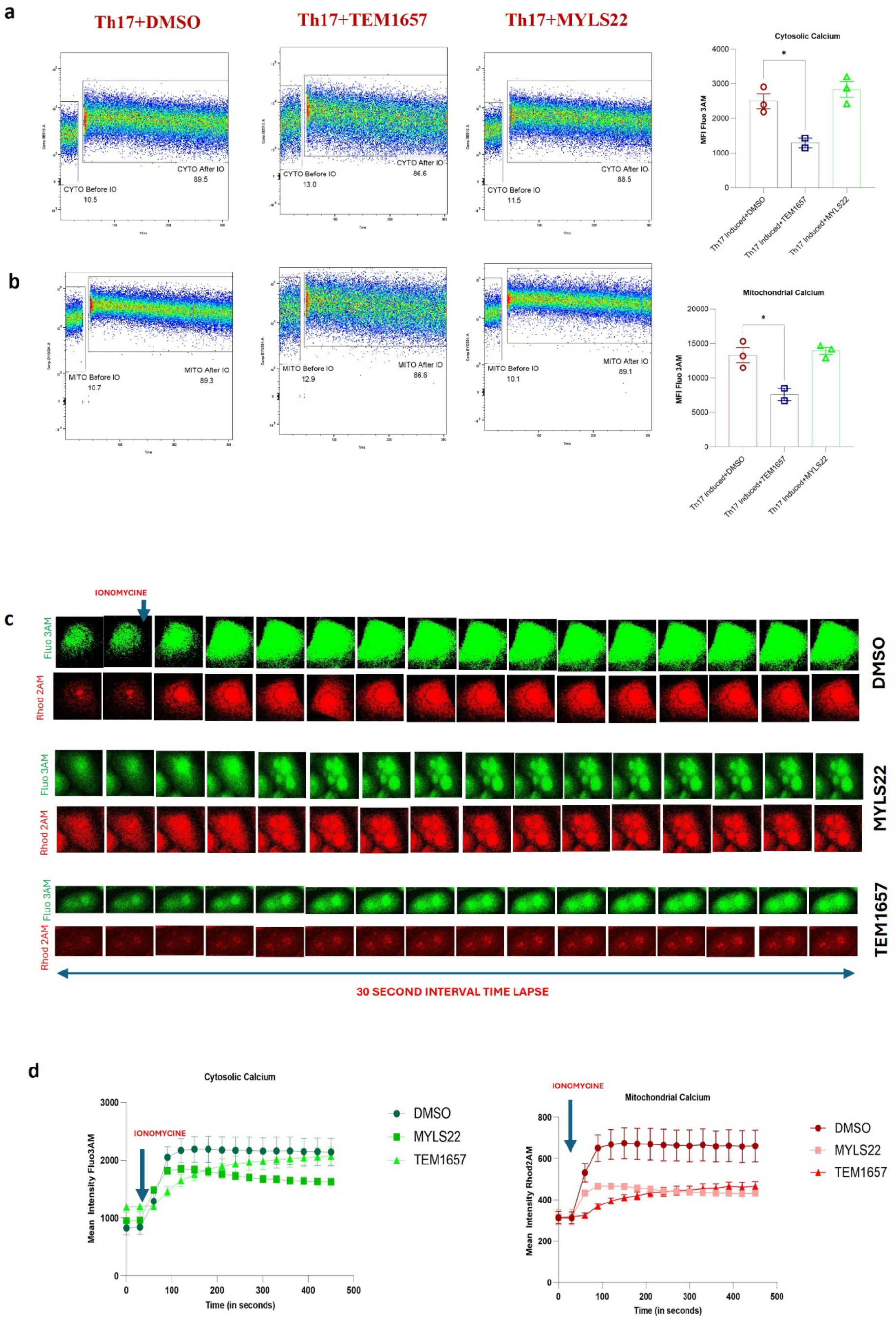
(a) Calcium influx analysis in cytosol and (b) mitochondria using flow cytometry after mobilizing calcium ions with ionomycin in the KCs (c) Live cell time-lapse confocal microscopic assessment of calcium migration from cytosol to mitochondria before and after ionomycin induction-the represented time-lapse series after 30 sec time intervals of cytosolic (green) and mitochondrial (red) calcium staining in KCs. (d) Graphical representation of cytosolic and mitochondrial calcium levels. The data are presented as the means ± standard errors of the means (S.E.M.) Statistical analysis was conducted by one way ANOVA and multiple comparisons were conducted with the help of Dunnett’s multiple comparisons test. *p < 0.05.

### TEM1657 impacts mitochondrial bioenergetics and T cell proliferation

As OPA1 is a critical regulator of, mitochondrial fusion, cristae organization, and oxidative phosphorylation; and previous reports showing beneficial effects of targeted OPA1 deficiency in inflammatory settings^41^, we speculated if TEM1657 has any impact on mitochondrial homeostasis. For this, real-time oxygen consumption rate (OCR) and extracellular acidification rate (ECAR) were measured in Th17-induced KCs in the presence and absence of TEM1657 with the help of Seahorse metabolic flux assay. Interestingly, we observed that both OCR and ECAR were consistently reduced in the presence of TEM1657 compared with DMSO control. Moreover, parameters including basal respiration, proton leak, maximal respiration, spare respiratory capacity, ATP-linked respiration, and non-mitochondrial OCR were also altered after only 90 minutes of TEM1657 exposure (Sup Fig. S3a). Similar results were obtained when KCs were co-stimulated with Th17 cytokines (Fig. 6a, b; Sup Fig. S3b). Our findings indicate that TEM1657 impairs oxidative phosphorylation in KCs, reducing overall energy demand. Importantly, the concomitant decrease in ECAR suggests that cells did not compensate by upregulating glycolysis but rather entered a metabolically quiescent state. Sustained reductions in OCR may reflect a loss of mitochondrial mass and increased fragmentation, as OPA1 inhibition prevents mitochondrial fusion. In addition, because OPA1 is essential for maintaining cristae structure, its suppression may contribute to disassembly of electron transport chain complexes, consistent with the reduced basal and maximal respiratory capacity observed in TEM1657-treated KCs.

**Figure 6.**
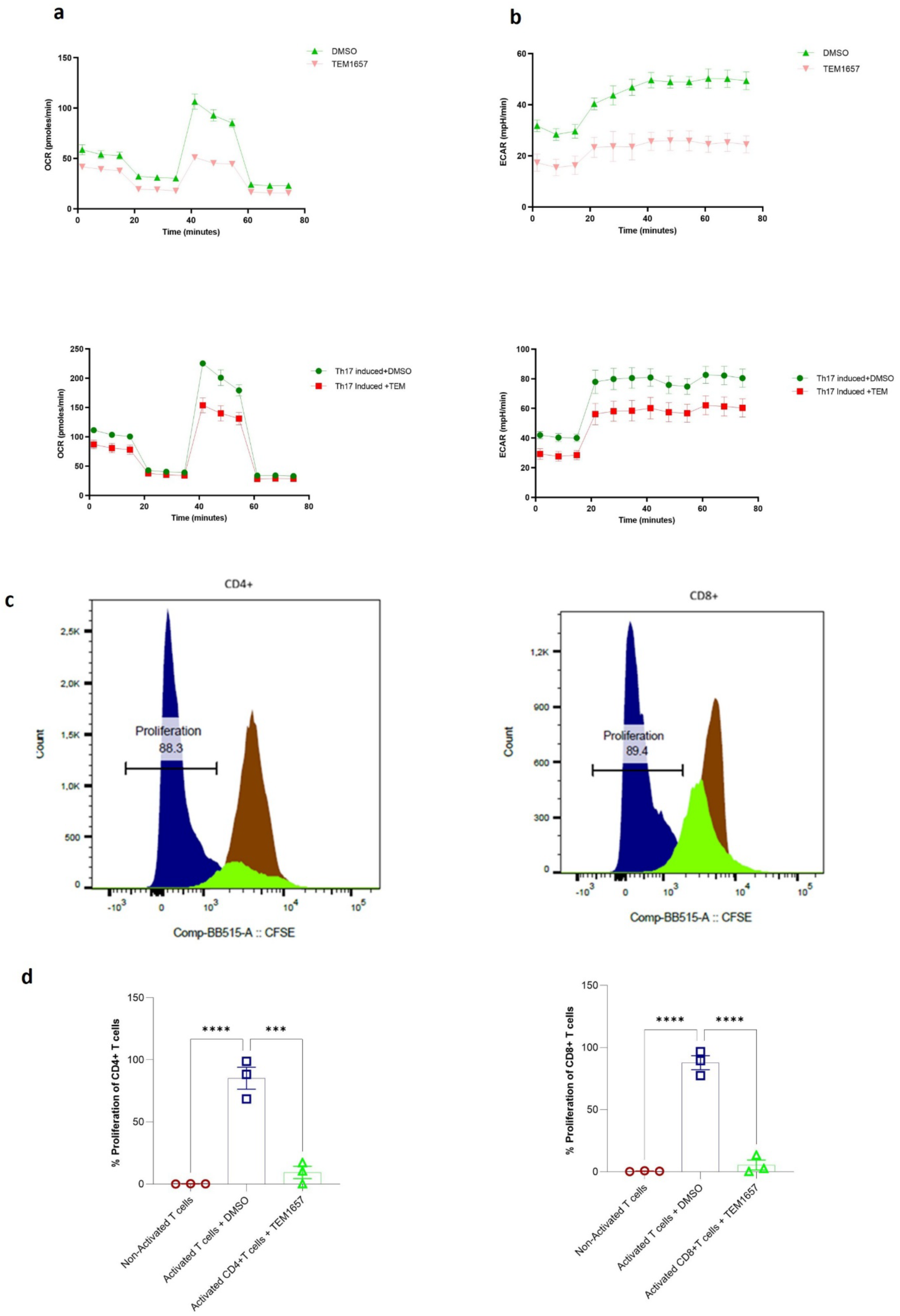
(a) Real-time oxygen consumption rates (OCR) and (b) real-time extracellular acidification rates (ECAR) of KCs after incubation with TEM1657 with and without Th17 stimulation. (c) CFSE-labelled T-cells non-activated (brown), activated with anti-CD3/CD28 and incubated with DMSO (blue) and activated with anti-CD3/CD28 and incubated with TEM1657 (green) stained with anti-CD4 and anti-CD8 antibodies and acquired on flow-cytometry. (d) Proliferation percentage of CD4+ and CD8+ T cells. The data are presented as the means ± standard errors of the means (S.E.M.) Statistical analysis was conducted by one way ANOVA and multiple comparisons were conducted with the help of Dunnett’s multiple comparisons test. ***p < 0.001, ****p < 0.0001.

**Figure 7.**
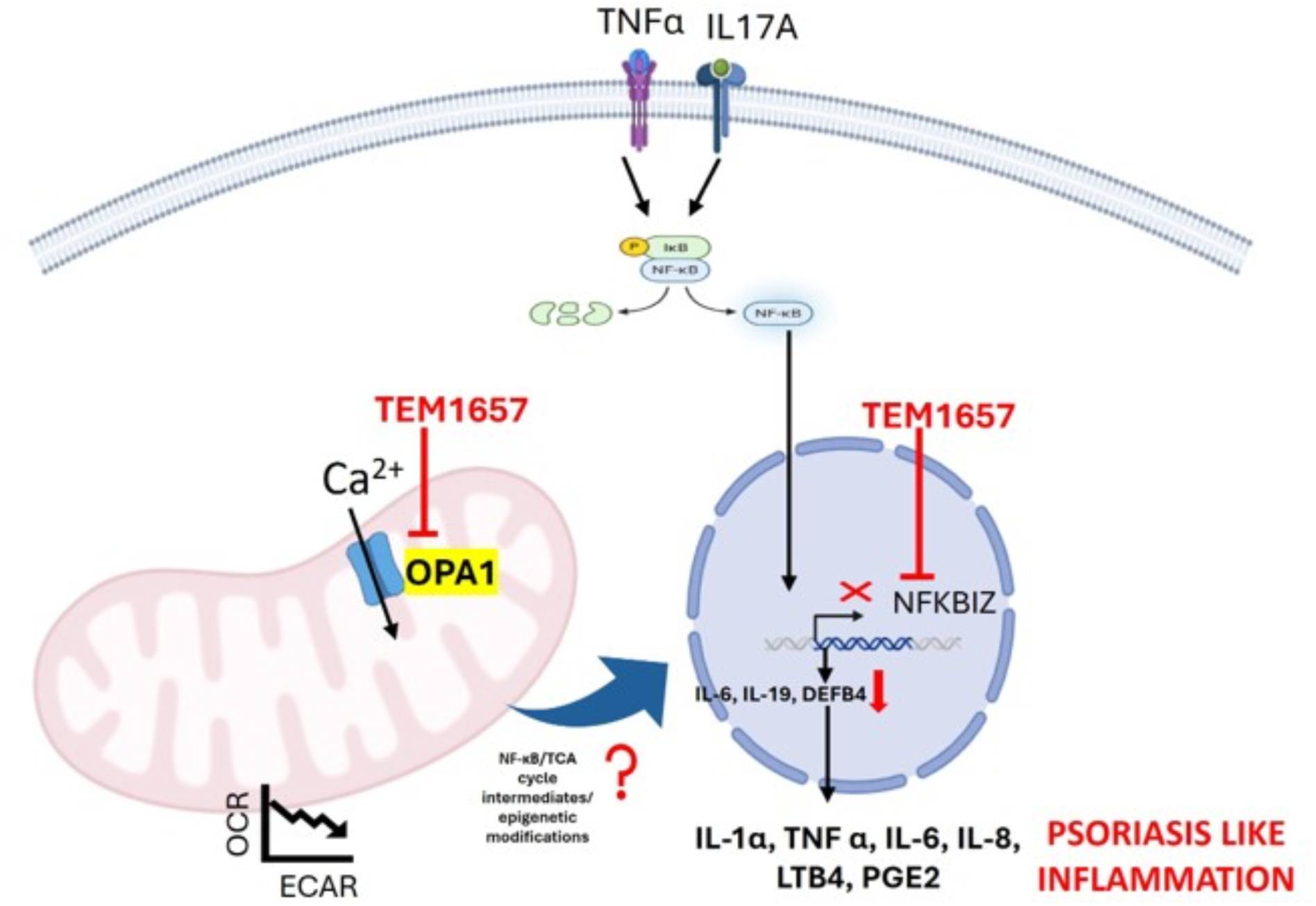
Schematic representation of potential targets and mechanism of action of TEM1657.

As psoriasis is also driven by abnormal T cell proliferation, we proceeded to TEM1657’s influence on CD3/CD28-induced proliferation of both CD4+ and CD8+ T lymphocytes. Notably, proliferation of CFSE-labelled CD4+ and CD8+ was observed to be significantly reduced in the presence of TEM1657(Fig. 6c, d). Certainly, the cumulative impact of TEM1657 on OPA1 and mitochondrial homeostasis might have rendered these immune cells in a metabolically scarce stage to progress rapid cell division after activation, thereby impacting the overall proliferation rate. Collectively, these results support the immunosuppressive potential of this molecule.

## Discussion

Although advances and evolution of biologics, together with the use of corticosteroids, have considerably improved the management of psoriasis, practical data suggest that their discontinuation often leads to frequent reoccurrence, making it the biggest challenge in psoriasis. The unmet need for treatment durability has driven increasing interest in precision medicine and the identification of new psoriasis-specific biomarkers^42,43^. Interestingly, in the past decade, NFKBIZ has emerged as a psoriasis-specific biomarker and therapeutic target. Localized suppression of NFKBIZ has been shown to attenuate IL-17-, IL-23-, and IL-36-driven inflammatory pathways, while offering a potential non-steroidal strategy that could circumvent the adverse effects associated with corticosteroid treatment^44–46^. Herein, we identify TEM1657, a novel small molecule that suppresses Th17-induced NFKBIZ expression through the modulation of OPA1-dependent mitochondrial homeostasis. To our knowledge, this is the exclusive signaling axis pathway targeted by TEM1657, which not only leads to suppression of imiquimod-induced psoriasis in mice but also inhibits NFKBIZ-dependent inflammatory response in KCs, thereby positioning TEM1657 as a novel candidate in the therapeutics of psoriasis.

In this study, we synthesized an analogue of 3,5-DiCQA, TEM1657, and tested its efficacy in an imiquimod-induced psoriasis-like mouse model. We then conducted its cellular uptake and subcellular distribution study in keratinocytes (KCs). Within 90 minutes of incubation, TEM1657 was found to accumulate in the chromatin-bound nuclear extract, demonstrating its ability to reach the nuclear compartment. To further explore its nuclear activity, we performed whole genome transcriptomics analysis in Th17-induced KCs. Notably, TEM1657 exhibited a broad anti-inflammatory transcriptional activity by suppressing the expression of several key pro-inflammatory cytokines and chemokines, as well as psoriasis-associated genes. We were exclusively driven by involvement of NFKBIZ in psoriasis due to the findings positioning it central to the therapeutics of psoriasis^26,47^. Herein, we demonstrated the ability of TEM1657 to suppress IκBζ/NFKBIZ expression and its downstream target genes. These observations suggest that TEM1657 targets a novel immunomodulatory mechanism with potential therapeutic relevance in psoriasis. Although additional studies are needed to establish precise downstream events in this mechanism of action, our data support a central role for NFKBIZ in the anti-inflammatory activity mediated by TEM1657.

We then further explored the molecular target of TEM1657 we performed CETSA® analysis and identified OPA1, MGLL, MICU1, and FECH as potential targets. Interestingly, we found that OPA1 inhibition recapitulated the suppression of NFKBIZ expression observed with TEM1657. These observations exposed a previously undiscovered link between OPA1 and NFKBIZ-dependent inflammatory signaling and support OPA1 as a candidate molecular target mediating the anti-inflammatory activity of TEM1657. Furthermore, as previous studies have identified NFKBIZ/IκBζ as a transcriptional co-activator of NF-κB, we therefore, decided to understand if TEM1657 influences activation of NF-κB. Importantly, our results show that this molecule does not affect NF-κB activation. These observations align with a recent study showing that inhibition of IκBζ/NFKBIZ does not affect the physiological activity of NF-κB^48,49^. This apparent selectivity may reflect the positioning of IκBζ downstream of NF-κB activation, where it regulates a specific subset of NF-κB-dependent inflammatory genes. Thus, TEM1657-mediated suppression of NFKBIZ may attenuate this inflammatory transcriptional program without interfering with upstream NF-κB activation^50^. Taken together, these results unveiled a potentially selective immunomodulatory mechanism where NFKBIZ-associated inflammatory profile in Th17-induced KCs can be attenuated without disturbing the crucial transcription factor NF-κB.

As OPA1 is involved in mitochondrial Ca^2+^ influx, and any alterations in cytosolic Ca2+ can impose changes in cellular-homeostasis. Additionally, Ca^2+^ also acts as a second messenger to various transcription factors including NF-κB, suggesting that any alterations in Ca^2+^ influx can significantly affect downstream targets of IL-17 pathway. We further demonstrate that TEM1657 significantly influences mitochondrial Ca^2+^ mobilization in Th17-induced KCs concomitantly with profound changes in mitochondrial-homeostasis. Taken together, these findings suggest that TEM1657 may act through OPA1 thereby altering cellular Ca^2+^ homeostasis.

Herein, we present the first experimental demonstration of a therapeutic compound that exhibits a distinct mechanism of action compared to current psoriasis therapies. Presently, the main line principal treatments includes biologics target cytokines IL17A, TNFα and 1L12/IL23^51^. Although these therapies are clinically active, their mechanisms primarily rely on the neutralization of specific extracellular cytokines or their signaling pathways. In contrast, TEM1657 exerts its anti-inflammatory effects through a distinct mechanism involving both OPA1-dependent mitochondrial homeostasis and NFKBIZ-dependent inflammatory signaling. Our findings provide proof of concept that modulation of mitochondrial homeostasis can be associated with suppression of the Th17-driven inflammatory response in psoriasis. However, a major limitation of our study is that, although we establish a functional link between OPA1 inhibition and NFKBIZ suppression, the molecular events connecting mitochondrial alterations to the regulation of NFKBIZ expression remain to be elucidated. Interestingly, a recent study showed that targeted deletion of OPA1 in Th17 cells leads to impaired Th17-effector cytokine production through the accumulation of TCA cycle intermediates, which further leads to epigenetic alterations^52^. Similarly, OPA1 loss in macrophages disrupts TCA cycle, further attenuating NF-κB signaling^41^. By contrast, in endothelial cells, OPA1 restrains NF-κB activity by controlling Ca^2+^ homeostasis, hence supporting angiogenesis^53^. Together, these observations highlight the cell-type specific role of OPA1 which fine-tunes inflammation in Th17 cells, macrophages and keratinocytes, making it potential intriguing target to shut down psoriasis-associated inflammation from multiple angles. Based on previous observations linking OPA1-dependent metabolic alterations to epigenetic regulation, we hypothesize that such metabolic changes could contribute to the suppression of NFKBIZ-dependent inflammatory responses. However, the involvement of TCA cycle intermediates and epigenetic mechanisms in TEM1657-mediated NFKBIZ regulation remains to be experimentally established.

Finally, we demonstrated that TEM1657 suppresses the proliferation of CD3/CD28 activated CD4+ and CD8+ T cells. As mitochondrial respiration is indispensable for cell-cycle progression and metabolic adaptations in activated T cells^54^. TEM1657-mediated modulation of OPA1 mitochondrial homeostasis may compromise the metabolic capacity to support T cell’s proliferation. These results are consistent with the recent study showing that OPA1 deficiency impairs the proliferation of rapidly dividing CD8+ effector T cells^55^. This new niche needs to be further explored in follow-up studies with OPA1 knock-out models together with comprehensive profiling of TCA cycle and epigenetic modifications. Moreover, the mechanisms linking OPA1-dependent mitochondrial alterations to NFKBIZ transcriptional regulation needs to be further elucidated. Such studies may clarify how these mitochondrial-modulations are translating into transcriptional expression of NFKBIZ and other psoriasis-associated genes.

In summary, this study identifies TEM1657 as a novel small molecule bearing therapeutic properties in IMQ-induced psoriasis mouse model. Our findings support a mechanism where TEM1657 modulates OPA1-dependent mitochondrial homeostasis in Th17-stimulated KCs, leading to a less metabolically active state and suppression of NFKBIZ-mediated inflammatory response. TEM1657-induced alterations in mitochondrial Ca²⁺ handling may contribute to the mechanistic link between mitochondrial dysfunction and transcriptional regulation, although the molecular events connecting these processes remain to be elucidated. In conclusion, TEM1657 acts at the intersection of OPA1-mediated mitochondrial homeostasis and NFKBIZ-dependent inflammatory signaling, unveiling a previously unrecognized functional axis in psoriasis-associated inflammation. These key findings pave the way towards novel therapeutic candidate which can be further explored to specifically target topical treatment of psoriasis.

## Materials and Methods

### (1R,2R,3R,5R)-2,5-dihydroxy-5-(octyl carbamoyl) cyclohexane-1,3-diyl (2*E*,2’*E*)-bis(3-(3,4-dihydroxyphenyl) acrylate) (TEM1657)

Unless otherwise specified, all compounds were obtained from commercial sources. The proton NMR was performed at 400 MHz using a Bruker BioSpin GmbH apparatus. Me_4_Si was the internal standard. The preparative-HPLC purification was performed on a column C18, 120 A, 10 µm, length: 250 mm, diameter: 50 mm.

3,5-Dicaffeoylquinic acid 3,5-DCQ (500 mg, 0.97mmol, 1eq) and 1-hydroxybenzotriazole HOBt (1.07 mmol, 144 mg, 1.1eq) were dissolved in dry DMF (2 mL) and anhydrous DCM (1 mL). 1-Ethyl-3-(3-dimethylaminopropyl) carbodiimide (EDCI, 1.07 mmol, 165 mg, 1.1 eq) was added dropwise at 0°C, then the mixture was stirred at the same temperature for 1 hour. Then octylamine (1.51 mmol, 195 mg, 250 mL, 1.5 eq) in DCM (0.5 mL) was added dropwise at 0°C. The mixture was stirred at 0°C for 30 minutes, then at room temperature for 2 hours. The reaction was stopped by adding HCl 1M (100 mL). Extraction was carried out using AcOEt (3*50 mL). The organic phase was washed successively with HCl 1M (2*50 mL) and brine (2*50 mL), then dried over MgSO_4_ *anhydrous*. The organic phase was concentrated to an oily yellow residue, which was dissolved in 4 mL of ACN and 4 mL of (H_2_O + 0,1% formic acid). Purification was carried out using reversed-phase column chromatography (ACN/water (+0,1% formic acid), gradient ACN: 45% to 50% in 25 min). We obtained 120 mg of product as a light-yellow solid (yield: 20%; purity: 98% UV).

^1^H NMR (400 MHz, MeOD) δ 7.63 (dd, *J* = 15.9, 6.5 Hz, 2H), 7.12 – 7.06 (m, 2H), 7.02 – 6.93 (m, 2H), 6.82 – 6.78 (m, 2H), 6.35 (dd, *J* = 24.6, 15.9 Hz, 2H), 5.55 (td, *J* = 10.4, 4.7 Hz, 1H), 5.48 (d, *J* = 3.2 Hz, 1H), 3.94 (dd, *J* = 9.6, 3.3 Hz, 1H), 3.19 (t, *J* = 6.5 Hz, 2H), 2.33 (dd, *J* = 15.3, 3.1 Hz, 1H), 2.18 (d, *J* = 9.0 Hz, 1H), 2.07 (t, *J* = 12.2 Hz, 2H), 1.50 (d, *J* = 6.5 Hz, 2H), 1.30 (s, 10H), 0.89 (t, *J* = 6.6 Hz, 3H).

^13^C NMR (100 MHz, MeOD) δ 177.6, 169.0, 168.9, 149.6, 149.5, 147.2,147.0, 146.9, 146.8, 127.9, 127.7, 123.0, 116.5, 115.7, 115.2, 115.1, 76.6, 73.8, 72.6, 71.7, 40.4, 39.6, 36.8, 32.9, 30.42, 30.3, 27.9, 23.7, 14.5.

MS (ESI) *m/z* calcd for C_33_H_42_NO_11_ [M+H]^+^ 628.28, found: 628.0.

### ELISA

For quantification of IL-8 and PGE2, KCs were stimulated with PMA (10ng/ml) and for IL-6 and TNFα, cells were stimulated with PMA (10ng/ml) + calcium ionophore A23187 (2 µM). LTB4 secretion was stimulated with the help of arachidonic acid (1 µM) + calcium ionophore A23187 (2 µM). Cells were pre-incubated with all the respective stimulants for 2 hours, followed by co-incubation with 3,5-DiCQA (50µM), TEM1657 (50µM) and positive control, Dexamethasone (10 µM) for IL-8, IL-6, TNFα and PGE2; NDGA (2µM) for LTB4 for 24 hours to quantify IL-8, PGE2, IL-6, TNFα and 2 hours for LTB4. Supernatants were collected after the respective incubation period and stored at −80°C until further use. Human IL-8/CXCL8 Quantikine ELISA Kit (R&D Systems, D8000C), Prostaglandin E2 ELISA Kit – Monoclonal (Cayman chemical company, 514010), Human IL-6 Quantikine ELISA Kit (R&D Systems, D6050), Human TNF-alpha Quantikine Kit (R&D Systems, DTA00C) and Leukotriene B4 ELISA Kit (Cayman chemical company, 520111) kits was used to quantify IL-8, PGE2, IL-6, TNFα and PGE2, respectively. The manufacturer’s protocol was followed and cytokine concentration as measured.

### Mice

Female Balb/c mice utilized in this study were purchased from CHARLES RIVER Breeding Center Laboratories, L’Arbresle, France. The housing conditions were employed such as mice were randomly distributed in cages in pairs, temperature of 24 ± 2°C, relative humidity of 50 ± 20%, 12-hour reverse light and dark cycle, M20 standard diet (SDS Dietex, Argenteuil, France) and unrestricted access to tap water was provided.

### Ethical approval

The animal protocols were performed after obtaining ethical approval from the expertise of the Lorraine Ethics Committee on Animal Experimentation (CELMEA). The animals were treated as per the guidelines of Directive 2010/63/EU of the European Parliament and of the Council of September 22, 2010 and Decree No. 2013-118 of February 1, 2013 on the protection of animals used for scientific purposes, as well as the ethical rules laid down by the ASAB (2012) and the Canadian Council on Animal Care (1993). All measures were included to reduce the pain of the animals.

### IMQ Psoriasis-like skin model and treatments

After 8 days of acclimatization period to the laboratory conditions, animals were weighed and randomly distributed into 7 groups (n=8) according to the treatment. Animals were shaved on the back three days prior, then psoriasis was induced by utilizing Aldara cream containing 5% imiquimod (Aldara™ 5% cream, Meda AB, Solna, Sweden, Lot no. GQL006A). Different testing concentrations of TEM1657 were prepared using Excipal (Excipial® Formula, Galderma International, la Défense, France, Lot T012). Dermoval cream (Dermoval® 0,05%, Laboratory GlaxoSmithKline, Marly-le-Roy, France, lot no. C760478) containing 0.05% clobetasol propionate was used for topical treatment of psoriasis. Psoriasis was induced by daily skin application for 10 days with 70 mg Aldara cream. Excipal containing 4 different concentrations of TEM1657 and Dermoval was administered cutaneously from D7 to D10.

### PASI score

Psoriasis area and severity index (PASI) score was calculated for assessing the disease progression. Erythema, degree of induration and degree of desquamation were studied and scored between 0 and 4 for none to very significant manifestation, respectively^56^. Also, the body weight was monitored 3 times a week, and the spleen weight was recorded at the end of the experiment.

### RT-qPCR

To determine the expression of NFKBIZ and IL-6, total RNA from cells was extracted with NucleoSpin RNA kit (MACHEREY-NAGEL, 740955) by following the manufacturer’s protocol. RNA concentration and quality were assessed using a NanoDrop spectrophotometer (Wilmington). Reverse transcription was performed with 500 ng RNA using ScriptTM cDNA synthesis kit (Bio-Rad). qPCR was performed with SYBR Green PCR master mix using QuantStudio 5 (Thermo Fisher Scientific) PCR machine. CT values were normalized to housekeeping genes, Ribosomal Protein L29 (RPL29). The sequence details of primers utilized are included in Supplementary Table S1. To determine the gene expression of IL-19 and DEFB4, total RNA was extracted with Qiagen RNeasy Kit (Qiagen, 74106) and cDNA was synthesized using the high-capacity RNA-to-cDNA kit (Applied Biosystems, 438706) by following the manufacturer’s recommendations. Finally, gene expression was assessed using TaqMan gene expression assays (Applied Biosystems) using probes Hs00604657_ml and Hs00823638_ml for IL-19 and DEFB4, respectively. The gene expression was normalized with the housekeeping gene beta-2-microglobulin and its expression was studied using probes Hs00984230_ml. Expression was assessed on Quantstudio 7 (Applied Biosystems).

### Western Blotting

Keratinocytes were lysed and proteins were separated using NuPAGE 10% Bis-Tris (Thermo Fisher Scientific, NP0342) gels and electroblotted for 1 hour on polyvinylidene fluoride (PVDF) membranes using Mini Blot Module (Thermo Fisher Scientific; B1000). Further, membranes were blocked using PBS solution containing 0.1% Tween (PBS-T) and 2% ECL blocking agent for 2 hours, followed by primary antibodies, Anti-Rabbit IκBζ (Cell Signaling Technology, 9244, dilution 1:1000) and Anti-mouse β-actin (Cell Signaling Technology, 3700, dilution 1:1000) incubation for 2 hours. Secondary antibodies Rabbit IgG HRP-linked whole Ab (GE Healthcare-Amersham, NA934) and Mouse IgG HRP-linked whole Ab (GE Healthcare-Amersham, NA931VS) were incubated at a dilution of 1:200,000 for 1 hour at room temperature. Proteins were detected using ECL detection reagent (Amersham, RPN2235). Protein bands were detected using G box Chemi XRQ (Syngene, 15472280).

### Cell culture

hTERT-immortalized Keratinocytes were obtained from ATCC (CRL-4048) and cultured following the manufacturer’s protocol. After sub-culturing, cells were treated with TNFα (2 ng/mL) and IL-17A (50 ng/mL) and TEM1657 (6.25 μM) for 90 minutes. In all *in vitro* experiments, DMSO was used as a control. After 90 minutes, cells were washed with PBS and stored at −80°C for RNA extraction. For microscopy and Seahorse experiments, cells were immediately stained or processed with respective protocols.

### PK/PD TEM1657 in whole blood sample and keratinocytes

Whole blood samples were incubated with TEM1657 (10 μM) for 0 min, 30 min, 1 hour, 3 hours, 6 hours, and 16 hours at 37°C. The blood sample was centrifuged at 2000 rpm for 10 min to separate plasma and blood cells. Further, 50 µl of sample (plasma/blood) was suspended in 450 µl of acetonitrile, and the samples were homogenized using sonication. The suspension was centrifuged at 13000 rpm for 5 minutes at 4°C to eliminate the precipitate. Finally, supernatants were diluted with an equal volume of DMSO/water, pH 3.0, 70/30 v/v and stored at −20°C for further analysis.

Keratinocytes were incubated with TNFα (2 ng/mL) and IL-17A (50 ng/mL) and TEM1657 (6.25 μM) for 10 minutes, 30 minutes and 90 minutes. Further, cytoplasmic extract, membrane extract, soluble nuclear extract and chromatin-bound nuclear extract were separated using the Subcellular Protein Fractionation Kit for Cultured Cells (78840, Thermo Scientific), and supernatant was stored at −20°C for further analysis.

### SRM

Supernatants from blood, plasma, and cellular compartment samples were thawed and diluted 50-fold with DMSO/EtOH 70/30 before analysis. SRM – single reaction monitoring – a method used for targeted quantification of compounds using MS/MS. The parent ion of the compound of interest (TEM1657, *m/z* 628, POS) is selected and transmitted by the first quadrupole (Q1, first selection stage) and fragmented in the second quadrupole (Q2). One of its fragment ions (in this case – 163, POS) is then filtered and transmitted through the third quadrupole (Q3, second selection stage) to the detector. Only compounds that meet these two selection stages (TEM1657 and its position isomers) are detected = high selectivity and sensitivity (due to limited noise).

### Transcriptomics

Keratinocytes were stimulated with TNFα (2 ng/mL) and IL-17A (50 ng/mL) and simultaneously treated with TEM1657 (6.25 μM) and DMSO for 90 minutes. Further, whole transcript Affymetrix microarrays and Affymetrix GeneChip^TM^ Human Clairom^TM^ S arrays (Thermo Fisher Scientific) were used to process RNA for whole genome transcriptomics analysis by following the manufacturer’s protocol. The normalized CEL files were analyzed using Transcriptomics analysis console 4.0 (Applied biosystems).

### TransAM

Keratinocytes were stimulated with TNFα (2 ng/mL) and IL-17A (50 ng/mL) and simultaneously treated with TEM1657 (6.25 μM) and DMSO for 6 hours. Nuclear proteins were prepared from the cells with the help of TransAM Nuclear Extract Kit (Active Motif, 40010) by following the manufacturer’s protocol. After protein quantification, 6 μg of nuclear extract was prepared for the NF-κB and STAT3 DNA binding assay. The activation of the p65 subunit of NF-κB and STAT3 was assessed by using TransAM^®^ NF-κB Activation Assays (Active Motif, 40096) and TransAM^®^ STAT3 (Active Motif, 45196). The manufacturer’s protocol was followed.

### CETSA®

Keratinocytes were incubated with TNFα (2 ng/mL) and IL-17A (50 ng/mL) for 15 minutes and then harvested with Trypsin/EDTA. Cells were then incubated with TEM1657 (10 μM) for 30 minutes at 37°C. After compound incubation, the sample was divided into 12 aliquots and a temperature range of 44°C to 66°C was applied. Cells were lysed with three cycles of freeze-thaw and samples from different temperatures were then pooled. The soluble fraction was separated with centrifugation at 30,000g for 20 minutes and further subjected to quantitative MS analysis. The experiment was performed at Pelago Bioscience (Sweden).

### Protein inhibition assay

Keratinocytes were stimulated with TNFα (2 ng/mL) and IL-17A (50 ng/mL) and simultaneously treated with TEM1657 (6.25 μM), DMSO, MGLL inhibitor JZL184 (10 μM), OPA1 inhibitor MYLS22 (50 μM), MICU1 inhibitor MCU i4 (10 μM), and FECH inhibitor NMPP (10 μM) for 90 minutes. Further, the cells were processed for RT-qPCR.

### Docking simulation

The crystal structures of the catalytic domain (PDB: 6JTG)^57^and dimer (PDB: 8EEW)^58^ were downloaded from the Protein Data Bank. TEM1657 conformer was prepared in 2D using ChemBioDraw Ultra 14.0 and transformed into 3D using VEGA 3.1.1^59^. The docking simulation was performed using AUTODOCK VINA 1.1.1. ^60,61^. The grid box was adjusted to comprise the complete surface of the protein. The analysis of binding mode and image generation was achieved using Pymol^62^ and ChimeraX 1.8.^63,64^

### Proliferation assay

T cells were obtained from the PBMCs of a healthy donor using Pan T cell isolation kit (Miltenyi, ref. 130-096-535) by following the manufacturer’s protocol. Further, activation of T lymphocytes was performed by mimicking CD3 and CD28 activation. For this, a round-bottomed 96-well cell culture plate was precoated with anti-CD3 antibodies (R&D Systems, ref: MAB100) and anti-CD28 antibodies (R&D Systems, ref: MAB342) were used at a concentration of 10 μg/mL for 4 hours. The purified T cells were labelled with carboxyfluorescein succinimidyl ester (CFSE, BD Biosciences, ref: 565082) at 0.02 mM for 10 minutes. Incubations were performed in RPMI 1640 with 10% fetal calf serum and 100 U/mL penicillin‒streptomycin (Life Technologies). CFSE-labelled T cells (5 x 10^4^ cells/well) were then added to the anti-CD3 and anti-CD28 precoated 96-well plate along with TEM1657(6.25 μM)/DMSO and incubated for 5 days. Finally, CD4+ and CD8+ T lymphocyte proliferation was analyzed by staining cells with the respective flow cytometry antibodies and cells were acquired on FACSymphony A3 (BD Biosciences), and analysis was performed on FlowJo V10.

### SeaHorse mitochondrial stress test

Keratinocytes were seeded in Seahorse XFp FluxPak (Agilent, 103022-100) at the concentration of 3X10^5^ cells/well. After 24 hours cells were stimulated with TNFα (2 ng/mL) and IL-17A (50 ng/mL) and simultaneously treated with TEM1657 (6.25 μM)/DMSO for 90 minutes. Further, Mito stress was assessed with the help of Seahorse XF Cell Mito Stress Test Kit (Agilent, 103010-100) by following the manufacturer’s protocol and the plates were acquired on Seahorse XF HS Mini Analyzer (Agilent).

### Confocal microscopy

Keratinocytes (2X10^4^ cells/well) were seeded in Nunc™ Lab-Tek™ II CC2™ Slide (Thermo) Culture Chamber System for 48 hours. The well adhered cells were then treated with TEM1657 (6.25 μM)/DMSO for 90 minutes. After 90 minutes, cells were washed with PBS and stained with Fluo-3 AM (5 μM) and Rhod-2AM (1 μM) for 30 minutes at 37°C. Cells were washed twice with PBS and kept at 37°C for an additional 20 minutes. Intracellular calcium was mobilized by exposing cells to 1 μM calcium ionomycin. Time-lapse confocal microscopy was performed on (Leica SP5 X AOBS LCSM, Leica microsystem, Wetzlar, Germany) using a 40X water objective. Fluorescence was sequentially excited to prevent spectral overlap with 488 nm (emission 514-550 nm) and 514 nm (emission564-606 nm) laser for Fluo-3 AM and Rhod-2AM, respectively. Acquisition of time-lapse images was performed at 1024 x 1024 pixels with a physical pixel size of 0.38μm x 0.38μm, and a single Z-plane per time point. Time-lapse was performed with intervals of 30s for 6-7 minutes. Images were acquired using Leica LAS AF and analyzed in ImageJ software.

### Flow cytometry for calcium influx

Keratinocytes were treated with TNFα (2 ng/mL) and IL-17A (50 ng/mL) and simultaneously treated with MYLS22 (50μM)/ TEM1657 (6.25 μM)/ DMSO for 90 minutes. Cells were then stained with Fluo-3 AM (5 μM) and Rhod-2AM (1 μM) for 30 minutes at 37°C. Further, cells were washed, trypsinized and passed through a Corning® 100µm Cell Strainer to obtain a homogeneous cell suspension. The acquisition was performed on FACSymphony A3 (BD Biosciences). After acquiring cells for 30 sec, cells were exposed to 1 μM calcium ionomycin. FCS files were analyzed with the help of FlowJo V10.

## Acknowledgements

The authors thanks Huguette Louis and Dominique Dumas from respectively, the cytometry facility and the confocal imaging facility of the SMP of Université de Lorraine. The authors gratefully acknowledge the financial support provided through the European Eurostars programme, within the framework of the PsoriaCure project. This project has also benefit from grant obtained by the region Grand-Est and FEDER.

## Conflict of interests

DM has received honoraria from Temisis therapeutics. Author Contributions (CRediT):

Conceptualization: P.G., D.M.

Methodology: P.G., A.B.A., G.J., L.B., J.Be., J. Br., A.E., R.P., J.H., M.M., N.d.I.

Investigation: P.G., A.B.A., G.J., L.B., J.Be., J. Br., A.E., R.P., J.H., M.M., N.d.I.

Formal analysis: P.G., A.S., A.B.A., G.J., P.H., J.-F.G.

Data curation: P.G., L.B., J.Be., J. Br, A.E., R.P., J.H., M.M., N.d.I.

Resources: P.H., J.-F.G, F.B., D.M.

Supervision: F.B., D.M.

Writing – original draft: P.G.

Writing – review & editing: P.G., A.S., A.B.A., G.J., L.B., J.Be., J. Br, A.E., R.P., J.H., M.M., N.d.I., P.H., J.-F.G., M.-T.R., J.-Y.J., F.B., D.M.

Project administration: P.G., D.M.

Funding acquisition: P.H., J.-F.G,., J.-Y.J., M.-T.R., F.B., D.M.

All authors reviewed and approved the final version of the manuscript.

