## Supplementary files for "The Mitochondria-Modulating Small Molecule TEM1657 Suppresses NFKBIZ Pathway and Resolves Th17-Driven Psoriatic Inflammation"

### SUPPLEMENTARY FIGURES

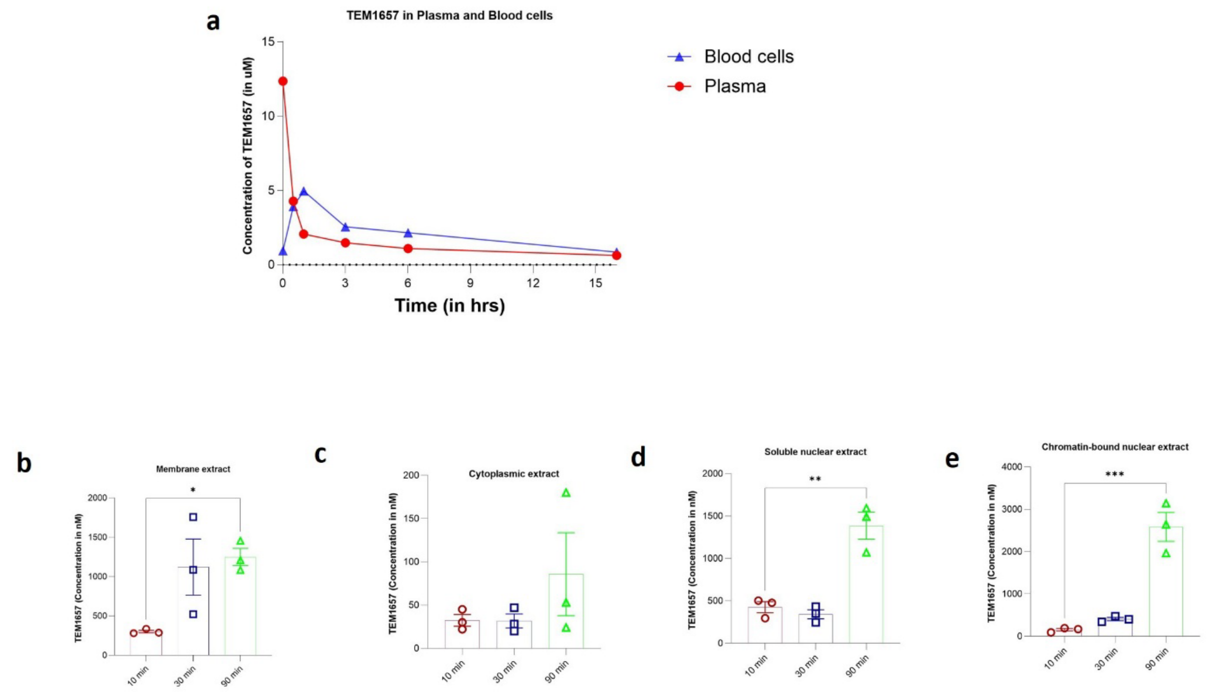


**Supplementary Figure S1.** PK/PD studies revealed that TEM1657 was cleared from plasma and could be distributed in blood cells within an hour; it also reached the keratinocyte nucleus within 90 minutes. Content in (a) blood cells and plasma sample measured at variable time points from 0-16 hours of incubation with TEM1657 (10µM); (b) membrane extract (c) cytoplasmic extract (d) Soluble nuclear extract (e) Chromatin-bound nuclear extract measured 10, 30 and 90 min of incubation with TEM1657 (6.25 µM). The data are presented as the means ± standard errors of the means (S.E.M.) Statistical analysis was conducted by one way ANOVA and multiple comparisons were conducted with the help of Dunnett's multiple comparisons test. *p < 0.05, **p < 0.01, ***p < 0.001.

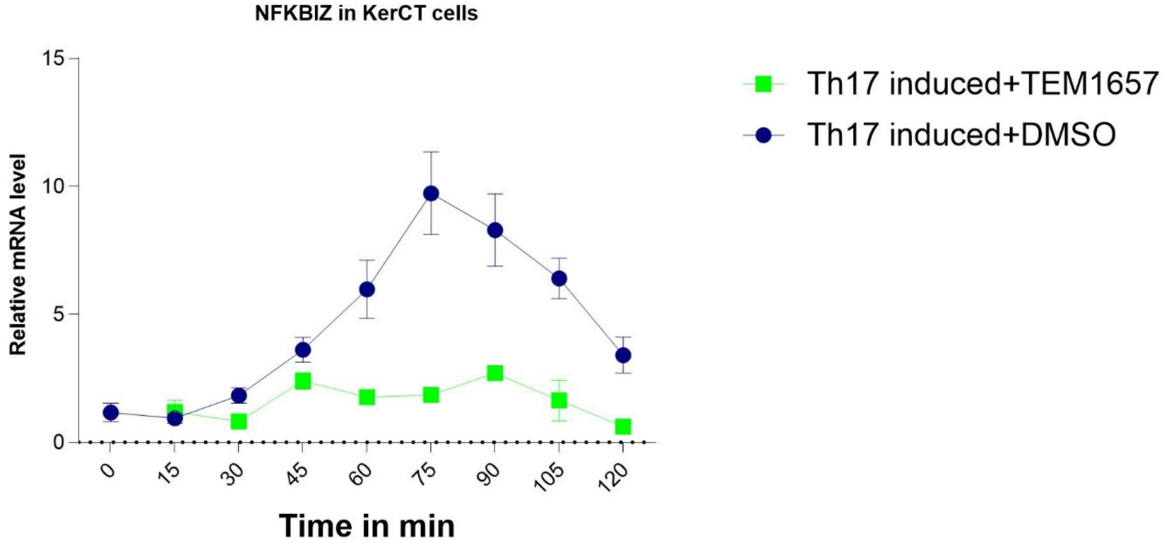


**Supplementary Figure S2.** Relative expression of NFKBIZ, tracked for 2hrs in KCs. Cells were induced and co-incubated with Th17 cytokines and TELM1657. RNA was collected at 9 time points (with an interval of 15 minutes) and expression of NFKBIZ was assessed with the help of RT-qPCR while normalizing with RP-29. The data are presented as the means ± standard errors of the means (S.E.M.).

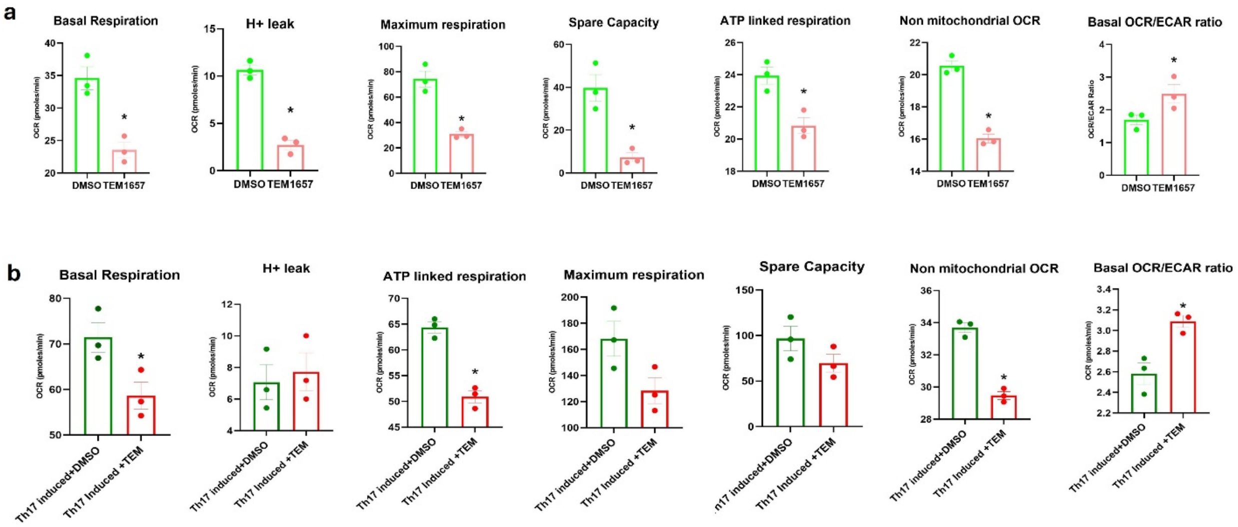


**Supplementary figure S3.** Mitochondrial respiratory functions in KCs treated with TEM1657. OCR and ECAR were used to calculate Basal respiration, H+ leak, maximum respiration, Spare capacity, ATP-linked respiration, non-mitochondrial OCR and Basal OCR/ECAR without Th17 induction **(a)** and with Th17 induction **(b)**. The data are presented as the means ± standard errors of the means (S.E.M.). n=3. The data are presented as the means ± standard errors of the means (S.E.M.) Statistical analyses were conducted by the Mann-Whitney test. *p < 0.05

**Supplementary table S1: List of primer sequences used for qPCR**

| Gene | Primer set sequence Forward 5’-3’ | Primer set sequence Reverse 5’-3’ |
|---|---|---|
| RP29 | GGGTCACCAGCAGCTGTACT | CCGATATCCTTCGCGTACTG |
| NFKBIZ | CCTGAGCTACTTCTACGGCG | ATATGGGGCTCAACTGGCTG |
| IL-6 | GGCTGAAAAAGATGGATGCT | GCTCTGGCTTGTTCCTCACT |

**Supplementary table S2: List of materials used**

| Name | Ref | Make |
|---|---|---|
| Rhod 2AM | R1245MP | Invitrogen |
| Ionomycin calcium salt | 124222 | |
| MYLS22 | #HY-136446/CS-0129450 | MedChemExpress |
| TNFa cytokine | 210TA-005 | R&D systems |
| IL-17A | 200-17 | Peprotech |
| RNA extraction kit | 740984 250 | Machese Nagel |
| N-methyl Protoporphyrin IX | 20846 | Cayman Chemical Company |
| MCU i4 | 39993 | Cayman Chemical Company |
| JZL184 | HY-15249 | MedChemExpress |
| Nordihydroguaiaretic acid (NDGA) | 74540 | Sigma |
| Dexamethasone | D4902 | Sigma |
| Phorbol 12-myristate 13-acetate (PMA) | 79346 | Sigma |
| Calcium Ionophore A23187 | C7522 | Sigma |
| Arachidonic acid | A3555 | Sigma |
| Human IL-8/CXCL8 Quantikine ELISA Kit | D8000C | R&D Systems |

| | | |
|---|---|---|
| Prostaglandin $E_2$ ELISA Kit - Monoclonal | 514010 | Cayman Chemical Company |
| Human IL-6 Quantikine ELISA Kit | D6050 | R&D Systems |
| Human TNF-alpha Quantikine Kit | DTA00C | R&D Systems |
| Leukotriene B4 ELISA Kit | 520111 | Cayman Chemical Company |
